# Chromosome-scale genome assembly of acerola (*Malpighia emarginata* DC.)

**DOI:** 10.1101/2024.07.30.605750

**Authors:** Kenta Shirasawa, Kazuhiko Harada, Noriaki Haramoto, Hitoshi Aoki, Shota Kammera, Masashi Yamamoto, Yu Nishizawa

## Abstract

Acerola (*Malpighia emarginata* DC.) is a tropical evergreen shrub that produces vitamin C-rich fruits. Increasing fruit nutrition is one of the main targets of acerola breeding programs. Genomic tools have been shown to accelerate plant breeding even in fruiting tree species, which generally have a long life cycle; however, the availability of genomic resources in acerola, so far, has been limited. In this study, as a first step toward developing an efficient breeding technology for acerola, we established a chromosome-scale genome assembly of acerola using high-fidelity long-read sequencing and genetic mapping. The resultant assembly comprises 10 chromosome-scale sequences that span a physical distance of 1,032.5 Mb and contain 35,892 predicted genes. Phylogenetic analysis of genome-wide SNPs in 60 acerola breeding materials revealed three distinct genetic groups. Overall, the genomic resource of acerola developed in this study, including its genome and gene sequences, genetic map, and phylogenetic relationship among breeding materials, will not only be useful for acerola breeding but will also facilitate genomic and genetic studies on acerola and related species.

## Introduction

Acerola (*Malpighia emarginata* DC.) is a tropical, fruit-bearing, evergreen shrub belonging to the family Malpighiaceae in the order Malpighiales. Acerola, also known as Barbados cherry and West Indian cherry, is native to Central and South America and the Caribbean Islands^1^. Acerola is cultivated in tropical and subtropical countries, especially in northwestern Brazil and the Mekong Delta region of Vietnam (Tien Giang Province and Ben Tre Province), primarily to meet the demand of the fresh fruit market and/or processing industries^2^. Because acerola fruits are rich in vitamin C (ascorbic acid), their concentrated juice is used as a raw material for preparing beverages and supplemental vitamin C powder. Therefore, one of the targets of acerola breeding programs is to increase the vitamin C content of acerola fruits. However, acerola, like other fruit tree species, has a long life cycle, making its breeding using conventional methods challenging.

Genomics can accelerate breeding programs not only in cereal and vegetable crops but also in fruit trees^3^. Indeed, novel genomics-based approaches, e.g., genome-wide association study and genomic selection based on machine learning methods, Bayesian networks, image analysis, and genomic prediction, have been implemented in the breeding programs of citrus, apple, and Japanese pear^4–6^. While massive genome information and multi-omics profiles have been available for the above-mentioned fruit trees^4,7^, the publicly available genome data of acerola, besides its chromosome number (2n = 20)^8^, has been limited. This situation has caused the breeding of acerola to lag behind that of other fruit crops.

High-fidelity long-read sequencing, also known as HiFi (PacBio) sequencing, has enabled genome assembly at the chromosome level or telomere-to-telomere level in several species^9^. The telomere-to-telomere assembly comprises a single contig, with telomere repeats at both ends, that covers the entire chromosome without any gaps of undetermined nucleotide sequences. Sequence scaffolding methods, together with the HiFi technology, help to establish chromosome-level genome assembly. Hi-C is a popular method used to connect the assembled sequences into the chromosome-level assembly^10^. Alternatively, genetic mapping with high-density genetic loci, based on a classical Mendelian law, enables anchoring the genome sequence fragments to linkage maps for establishing chromosome-level sequences^11^. The establishment of genome resources and genetic diversity assessment of breeding materials would be the first step in the breeding of acerola. In this study, we used HiFi technology to sequence the acerola genome and then anchored the sequences to a genetic map to construct a chromosome-level genome assembly of acerola. Subsequently, we assessed the genetic diversity of acerola breeding materials using SNPs derived by double digest restriction-site associated sequencing (ddRAD-Seq). The acerola genome information generated in this study will be helpful in accelerating acerola breeding programs and increasing knowledge of the genetics and genomics of Malpighiales.

## Materials and Methods

### Plant materials

Acerola cultivar NRA309, which was registered in 2008 by Nichirei Foods Inc. (Tokyo, Japan), was used for genome sequencing. An S1 mapping population (nC=C118), derived by the self-pollination of NRA309, was used to construct a genetic map of acerola (Supplementary Table S1). In addition to NRA309, a total of 59 acerola lines were used for genetic diversity analysis. These 59 lines included 10 lines from Brazil, four from Japan, and nine from USA as well as 36 progenies generated in breeding programs by crossing the above-mentioned lines (Supplementary Table S2). All plant materials were maintained at the Faculty of Agriculture, Kagoshima University.

### Genome size estimation

Genomic DNA (gDNA) of NRA309 was extracted from leaves using Genomic Tip (Qiagen, Hilden, Germany). The PCR-free Swift 2S Turbo Flexible DNA Library Kit (Swift Sciences, Ann Arbor, MI, USA) was used to construct a short-read library, which was then converted into a DNA nanoball sequencing library using the MGI Easy Universal Library Conversion Kit (MGI Tech, Shenzhen, China). The library was sequenced using a DNBSEQ G400 instrument (MGI Tech) to generate 100-bp paired-end reads. The genome size of NRA309 was estimated using Jellyfish (*k*-mer size = 17)^12^.

### Genome sequencing and primary assembly

NRA309 gDNA was subjected to long-read library preparation. Briefly, the NRA309 gDNA was sheared to an average fragment size of 40 kb using Megaruptor 2 (Deagenode, Liege, Belgium) in the Large Fragment Hydropore mode, and the sheared DNA was used for HiFi SMRTbell library preparation with the SMRTbell Express Template Prep Kit 2.0 (PacBio, Menlo Park, CA, USA). The obtained DNA libraries were fractionated with BluePippin (Sage Science, Beverly, MA, USA) to eliminate fragments shorter than 20 kb in length. The fractionated DNA libraries were sequenced on a SMRT Cell 8M on the Sequel II system (PacBio). HiFi reads were constructed using the CCS pipeline (https://ccs.how) and assembled using Hifiasm (version 0.16.1)^13^ with default parameters. Organelle genome sequences, identified by sequence similarity searches of the reported plastid and mitochondrial genome sequences of acerola relatives (Supplementary Table S3) using Minimap2 (version 2.24)^14^, were eliminated. Assembly completeness was evaluated with the embryophyta_odb10 data using Benchmarking Universal Single-Copy Orthologs (BUSCO) (version 5.2.2)^15^.

### Chromosome-level genome assembly by genetic mapping

The gDNA of lines in the S1 population was extracted from leaves using the Plant Genomic DNA Extraction Mini Kit (Favorgen, Ping-Tung, Taiwan). The extracted gDNA was digested with *Pst*I and *Msp*I restriction endonucleases and subjected to ddRAD-Seq library preparation^16^. The resultant library was sequenced on a DNBSEQ G400 instrument (MGI Tech) to generate 100-bp paired-end reads. After removing adapter sequences (AGATCGGAAGAGC) with fastx_clipper in the FASTX-Toolkit (version 0.0.14, http://hannonlab.cshl.edu/fastx_toolkit) and trimming low-quality reads (quality score < 10) with PRINSEQ (version 0.20.4)^17^, high-quality reads were aligned onto the genome assembly of NRA309 using Bowtie2 (version 2.3.5.1)^18^. High-confidence biallelic SNPs were identified using the mpileup and call options of BCFtools (version 1.9)^19^ and filtered using VCFtools (version 0.1.16)^20^ based on the following criteria: read depth ≥ 5; SNP quality = 999; minor allele frequency ≥ 0.2; proportion of missing data < 20%. A linkage analysis of the SNPs was performed using Lep-Map3 (version 0.2)^21^ to construct a genetic map. Contig sequences were anchored to the genetic map, and chromosome-level pseudomolecule sequences were established using ALLMAPS (version 0.7.3)^11^. Telomere sequences containing repeats of a 7-bp motif (5C-TTTAGGG-3C) were searched using the search subcommand of tidk (https://github.com/tolkit/telomeric-identifier), with a window size of 100,000 bp.

### Prediction of protein-coding genes and repetitive sequences

Protein-coding genes were predicted using the *ab initio* strategy of BRAKER3 (version 3.0.7)^22^ and Helixer (version 0.3.2)^23^. Prediction completeness was evaluated with the embryophyta_odb10 data using BUSCO (version 5.2.2)^15^. The predicted genes were functionally annotated using emapper (version 2.1.12) implemented in EggNOG^24^, in conjunction with searches against the UniProtKB database^25^ and Arabidopsis peptide sequences (Araport11)^26^ using DIAMOND (version 2.0.14)^27^ and BLAST^28^, respectively. Repetitive sequences in the assembly were identified using RepeatMasker (https://www.repeatmasker.org), based on repeat sequences registered in Repbase and a *de novo* repeat library built with RepeatModeler (https://www.repeatmasker.org).

### Genetic diversity analysis

The genome structure of acerola was compared with that of three Malpighiales species, including rubber tree (*Hevea brasiliensis*, NCBI RefSeq assembly GCF_030052815.1)^29^, cassava (*Manihot esculenta*, GCF_001659605.2)^30^, and castor bean (*Ricinus communis*, GCF_019578655.1)^31^, using Minimap2^14^ implemented in D-Genies^32^. ddRAD-Seq libraries of all 59 acerola lines were prepared as described above and sequenced on a DNBSEQ G400 instrument (MGI Tech). Subsequently, the ddRAD-seq reads obtained from all 59 lines and NRA309 were aligned onto the chromosome-level pseudomolecule sequences to detect sequence variants. High-confidence biallelic SNPs were selected with the filtering conditions: read depth ≥ 5; SNP quality = 999; minor allele frequency ≥ 0.05; proportion of missing data < 20%. The population structure of 60 lines was evaluated based on the maximum-likelihood estimation of individual ancestries using ADMIXTURE (version 1.3.0)^33^. A phylogenetic tree, based on 100 bootstrap replicates, was created with SNPhylo (version 20140701)^34^ and visualized with iTOL (version 6.9.1)^35^.

## Results

### Genome sequencing and assembly

To estimate the acerola genome size, clean short-read data (73.1 Gb) were subjected to *k*-mer distribution analysis. The results indicated that the acerola genome was highly heterozygous, with a haploid genome size of 1,114.2CMb (Supplementary Figure S1). A total of 90.3 million HiFi reads, with an N50 length of 32.4 kb (28.3 Gb; 24.6× coverage of the estimated genome size), were obtained from one SMRT Cell 8M and assembled into 73 contigs. Three potential contaminant sequences (221.1 kb in total) from organelle genomes were removed. Consequently, 70 contigs with a total length of 1,053.0 Mb and N50 and N90 lengths of 73.1 Mb and 39.1 Mb, respectively (Table 1), were obtained. These 70 contigs represented the genome assembly of acerola and were designated as NRA309_r1.0. A genetic map of acerola was constructed based on 940.0 million ddRAD-Seq reads obtained from 118 S1 lines and NRA309 (parental line). Briefly, high-quality reads were aligned to the assembled contigs as a reference, with an average mapping rate of 88.6% (Supplementary Table S1), and 7,935 high-confidence SNPs were detected. In the subsequent linkage analysis, 10 linkage groups were obtained, with the number of linkage groups corresponding to the basic chromosome number of acerola. The resultant genetic map spanned a distance of 1,059.8 cM and contained a total of 7,775 SNPs (Table 2, Supplementary Figure S2). Contigs were aligned onto the acerola genetic map to establish chromosome-level sequences. Of the 70 contigs, 19 were mapped to 10 linkage groups. Among them, 1, 2, and 3 contigs were mapped on 3, 5, and 3 linkage groups, respectively (Table 2). The 10 pseudomolecule sequences spanned a physical distance of 1,032.5 Mb (98.1% of the total contig length) (Tables 1, 2, Supplementary Figure S2). Telomere repeats were found at one end of seven sequences and at both ends of two sequences, while no telomeres were found in one sequence (Table 2). Overall, chromosomes 5 and 7 were completely assembled, without gaps, at the telomere-to-telomere level. The remaining 51 contigs (total length = 20.4 Mb) with five telomere repeat units were not assigned to any linkage groups (Table 1, 2). In accordance with the BUSCO score, the acerola genome assembly was 98.8% complete (Table 3).

**Table 1.** Statistics of the acerola genome assembly.

|  | Contig sequences<br>(NRA309_r1.0) | Pseudomolecule<br>sequences<br>(NRA309_r1.0.pmol) | Contigs unassigned to<br>any linkage groups |
| --- | --- | --- | --- |
| Total size (bp) | 1,052,960,172 | 1,032,546,981 | 20,414,091 |
| No. of sequences | 70 | 10 | 51 |
| N50 length (bp) | 73,099,088 | 104,183,651 | 1,133,687 |
| N90 length (bp) | 39,070,837 | 92,689,787 | 135,127 |
| Gap length (bp) | 0 | 900 | 0 |
| No. of genes | Not analyzed | 35,551 | 341 |

**Table 2.** Statistics of the acerola pseudomolecule sequence.

| Linkage group | No. of<br>SNPs | Map length<br>(cM) | Total length<br>(bp) | No. of<br>contigs | Telomere<br>repeat unit <sup>a</sup> | No. of<br>genes |
| --- | --- | --- | --- | --- | --- | --- |
| NRA309ch01 | 1,035 | 105.3 | 104,183,651 | 2 | 1 (Bottom) | 3,580 |
| NRA309ch02 | 982 | 126.3 | 118,493,997 | 3 | 0 (None) | 4,537 |
| NRA309ch03 | 906 | 100.5 | 92,689,787 | 3 | 1 (Bottom) | 3,215 |
| NRA309ch04 | 847 | 110.2 | 107,707,034 | 2 | 1 (Top) | 3,815 |
| NRA309ch05 | 781 | 87.9 | 100,396,053 | 1 | 2 (Both) | 3,540 |
| NRA309ch06 | 770 | 121.8 | 110,748,733 | 2 | 1 (Bottom) | 3,723 |
| NRA309ch07 | 748 | 125.6 | 122,825,995 | 1 | 2 (Both) | 4,097 |
| NRA309ch08 | 605 | 89.7 | 93,730,031 | 1 | 1 (Bottom) | 3,103 |
| NRA309ch09 | 562 | 90.4 | 81,726,457 | 2 | 1 (Bottom) | 2,593 |
| NRA309ch10 | 539 | 102.1 | 100,045,243 | 2 | 1 (Top) | 3,348 |
| Subtotal (ch01 to<br>ch10) | 7,775 | 1,059.8 | 1,032,546,981 | 19 | 11 | 35,551 |
| Unassigned<br>contigs | Not<br>available | Not<br>available | 20,414,091 | 51 | 5 | 341 |
| **Total** | **7,775** | **1,059.8** | **1,052,961,072** | **70** | **16** | **35,892** |

<sup>a</sup> Locations of telomere repeat units are shown in parentheses.

**Table 3.** Completeness of the genome assembly and predicted genes of acerola.

|  | Genome<br>assembly | Predicted<br>genes |
| --- | --- | --- |
| Complete | 98.8% | 99.1% |
| Single-copy complete | 95.6% | 95.8% |
| Duplicated complete | 3.2% | 3.3% |
| Fragmented | 0.6% | 0.7% |
| Missing | 0.6% | 0.2% |

### Gene and repetitive sequence prediction

Next, we predicted protein-coding genes in the acerola genome. First, a total of 196,853 genes, with an average coding sequence length of 1,617 bp, were predicted using BRAKER3. However, the BUSCO score of these genes was only 82.5% (41.1% single-copy complete BUSCOs and 41.1% duplicated complete BUSCOs). Because of the high gene number and low BUSCO score, we speculated that BRAKER3-based prediction was inaccurate. Therefore, we repeated the gene prediction using Helixer. As a result, a total of 35,892 genes (average coding sequence length = 1,150 bp) were predicted, of which 35,551 and 341 genes were located on the 10 chromosome-scale sequences and the 51 unassigned contigs, respectively. The BUSCO score of Helixer-predicted genes was 99.1% (Table 3). Out of the 35,892 genes, 31,742 (88.4%), 30,989 (86.3%), and 29,068 (81.0%) genes were functionally annotated with EggNOG mapper, DIAMOND search against UniProtKB, and BLAST search against Araport11 peptide sequences, respectively (Supplementary Table S4). In total, 32,239 (89.8%) genes were annotated by at least one of the three methods.

Repetitive sequences occupied a total of 816.3CMb (77.5%) of the pseudomolecule sequences. Nine major types of repeats were identified in varying proportions (Table 4). LTR retroelements (427.2CMb) represented the dominant repeat type in the pseudomolecule sequences, followed by DNA transposons (89.8CMb). Repeat sequences unavailable in public databases totaled 252.5CMb.

**Table 4.** Repetitive sequences in the acerola genome.

| Repeat type | No. of repeats | Total length (bp) | Relative proportion (%) |
| --- | --- | --- | --- |
| SINEs | 829 | 109,559 | >0.0 |
| LINEs | 51,728 | 30,364,578 | 2.9 |
| LTR elements | 372,431 | 427,204,241 | 40.6 |
| DNA transposons | 145,807 | 89,786,642 | 8.5 |
| Small RNA | 4,891 | 7,793,074 | 0.7 |
| Satellites | 1,995 | 437,534 | >0.0 |
| Simple repeats | 154,212 | 6,367,487 | 0.6 |
| Low complexity | 33,617 | 1,711,063 | 0.2 |
| Unclassified | 687,125 | 252,478,999 | 24 |

### Comparative genome structure analysis

The pseudomolecule sequences of acerola were aligned against those of rubber tree (*H. brasiliensis*), cassava (*M. esculenta*), and castor bean (*R. communis*), all of which are members of Malpighiaceae. Unexpectedly, no clear similarity was detected, and probable conserved structures were highly fragmented between the genome of acerola and those of the other three Malpighiaceae species (Supplementary Figure S3).

### Genetic diversity of acerola lines

To evaluate the genetic diversity of acerola lines, the gDNA of 59 lines was subjected to ddRAD-Seq. An average of 12.7 million ddRAD-Seq reads per sample were obtained. High-quality reads from 59 lines and NRA309 were aligned to the pseudomolecule sequences of NRA309 (reference), with an average mapping rate of 90.6% (Supplementary Table S2), and 49,070 high-confidence SNPs were detected. According to SnpEff results, the most prominent SNP type was modifier impact (59.5%) in introns and intergenic regions, followed by low impact (21.9%; synonymous mutations), moderate impact (18.2%; missense mutations), and high impact (0.4%; nonsense and splice-site mutations) (Supplementary Table S5). ADMIXTURE analysis grouped the 60 lines into three clusters (K = 3): cluster a, seven Japanese and Hawaiian lines; cluster b, 16 Hawaiian and Brazilian lines; and cluster c, 36 lines derived from crosses between FB05 and five members of cluster b (Figure 1a). These three clusters were well-supported by phylogenetic analysis (Figure 1b).

**Figure 1.**
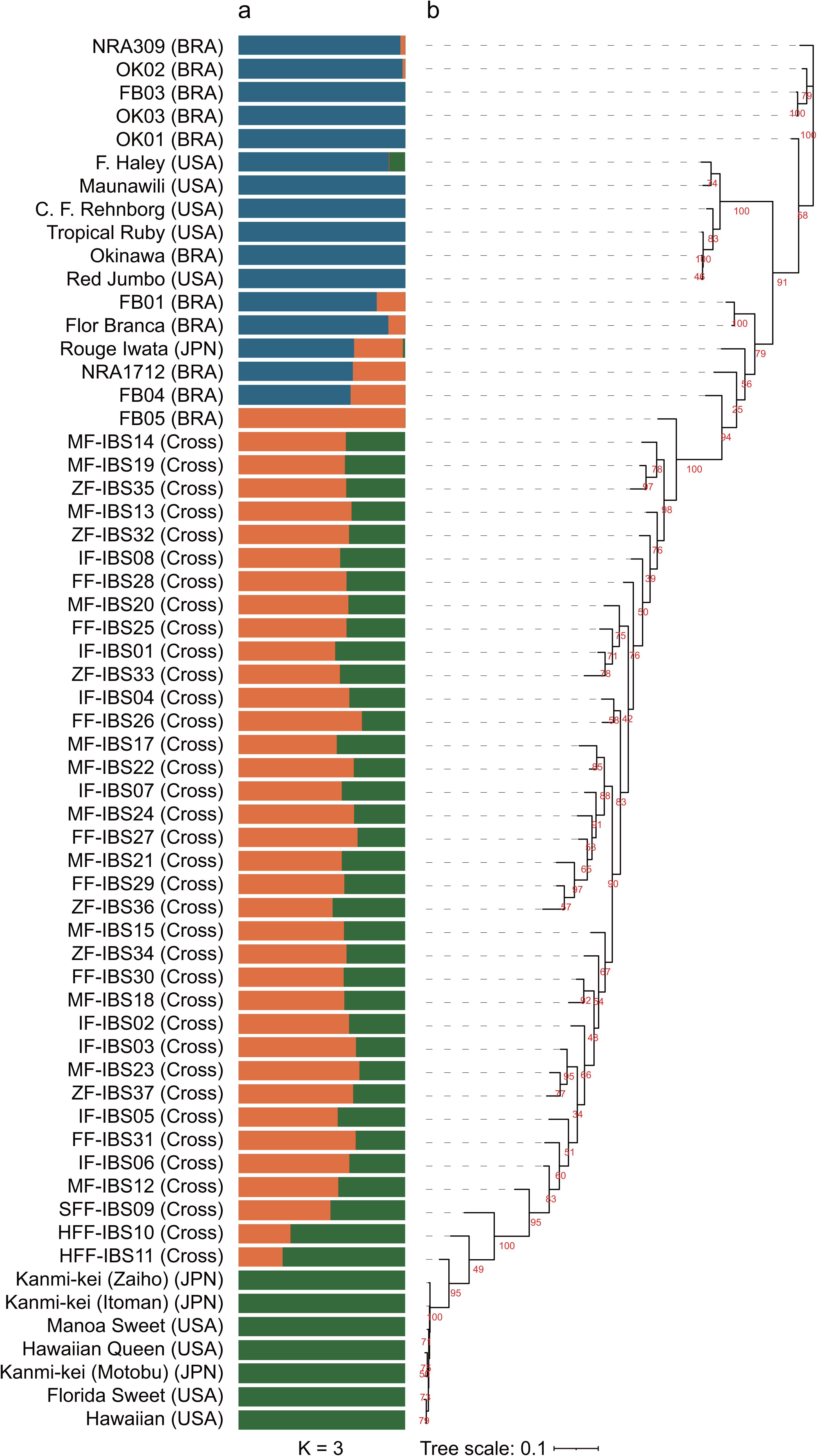
Genetic structure and phylogenetic tree of 60 acerola lines. (a) Population structure of 60 acerola lines. Each color represents a distinct group. The origin of acerola lines is shown in parentheses: BRA, Brazil; JPN, Japan; USA, United States of America; and Cross, Progenies of crosses. (b) Phylogenetic tree. Numbers in red on branches indicate bootstrap values based on 100 replicates.

## Discussion

Here, we present a chromosome-scale genome assembly of acerola, a member of the family Malpighiaceae. To the best of our knowledge, this is the first report of a chromosome-scale genome assembly of not only acerola but also a Malpighiaceae species. Chromosome-level genome assemblies have been reported for at least three species in the order Malpighiales, namely, rubber tree (*H. brasiliensis*), cassava (*M. esculenta*), and castor bean (*R. communis*), all of which are members of the family Euphorbiaceae. Comparative genome structure analysis revealed that the genome structures of acerola and the three Euphorbiaceae species were seldom conserved (Supplementary Figure S3). This suggests the possibility that acerola diverged from members of Euphorbiaceae within the order Malpighiales. More chromosome-level genome assemblies for members of the Malpighiaceae would be required to validate this hypothesis.

The acerola genome information generated in this study would be useful for acerola breeding programs, in which the improvement of fruit vitamin C content together with plant disease and pest resistance are frequently targeted. The phylogenetic relationship of acerola cultivars determined in this study (Figure 1) was in agreement with that determined previously using PCR-based sequence-related amplified polymorphism (SRAP) markers^36^. Parental lines in breeding programs could be adequately selected to enhance the genetic diversity of breeding materials. Furthermore, agronomically important loci could be identified through quantitative trait locus analysis, based on the genetic map. Once the genetic loci of interest have been determined, the underlying candidate genes could be easily identified using genome information. Being a woody plant, the life cycle or generation time of acerola is longer than that of annual herbaceous plants including vegetables, which makes the completion of conventional genetic analysis and breeding within a limited time period challenging. Therefore, the application of novel genomics-based approaches, e.g., genome-wide association study and genomic selection based on machine learning methods, Bayesian networks, image analysis, and genomic prediction^4–6^, has been proposed for fruit tree crops including woody plants.

We conclude that the acerola genome resources developed in this study, including its genome and gene sequences, genetic map, and phylogenetic relationship among breeding materials, would be useful for genomics research on acerola and Malpighiales and for the development of novel fast-paced breeding technologies. This study provides a standard genome resource for the genomics, genetics, and breeding of acerola and related species.

## Data Availability

Raw sequence reads were deposited in the DNA Data Bank of Japan (DDBJ) BioProject database under the accession number PRJDB18209, for which details are listed in Supplementary Tables S1 and S2. The assembled sequences are available at DDBJ (accession numbers: AP035802 - AP035862) and Plant GARDEN (https://plantgarden.jp)^37^.

## Acknowledgments

We thank Y. Kishida, C. Minami, K. Ozawa, H. Tsuruoka, and A. Watanabe (Kazusa DNA Research Institute) for technical assistance.

---

**Supplementary Figure S1** Estimated size of the acerola genome, based on *k*-mer analysis (*k* = 17), with the given multiplicity values.

**Supplementary Figure S2** Genetic and physical maps of the acerola genome. Left: SNP loci on the genetic map (vertical lines) and physical map (bars) are connected with horizontal lines. Right: Positions of SNP loci are indicated with dots on the genetic map (y-axis, cM) and physical map (x-axis, Mb).

**Supplementary Figure S3** Comparative analysis of the genome sequence and structure of acerola and three other Malpighiales species. Dots indicate similarities in the genome structures of rubber tree (*Hevea brasiliensis*), cassava (*Manihot esculenta*), and castor bean (*Ricinus communis*). Chromosome numbers are indicated above the x-axis and on the right side of the y-axis, and genome sizes (Mb) are shown below the x-axis and on the left side of the y-axis.

