## Supplementary Figure for "Chromosome-scale genome assembly of acerola (*Malpighia emarginata* DC.)"

**Supplementary Table S1** Number of ddRAD-Seq reads, map rate on the reference sequence, and accession numbers of ddRAD-Seq for an S1 mapping population via self-pollination of NRA309.
**Supplementary Table S2** Number of ddRAD-Seq reads, map rate on the reference sequence, and accession numbers of breeding materials.
**Supplementary Table S3** Reference genomes used for removing organelle sequences from the acerola genome assembly.
**Supplementary Table S4** Functinal annotation for the predicted genes in the acerola genome.
**Supplementary Table S5** Annotation of variants detected among 60 acerola lines.

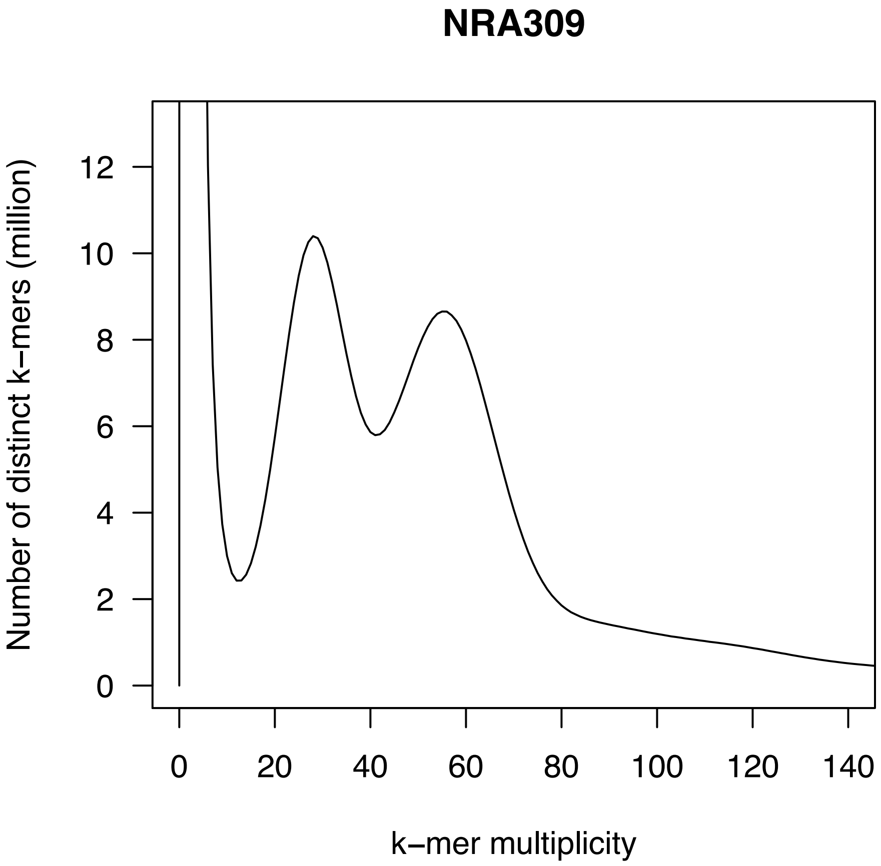


**Supplementary Figure S1** Estimated size of the acerola genome, based on *k*-mer analysis ($k = 17$), with the given multiplicity values.

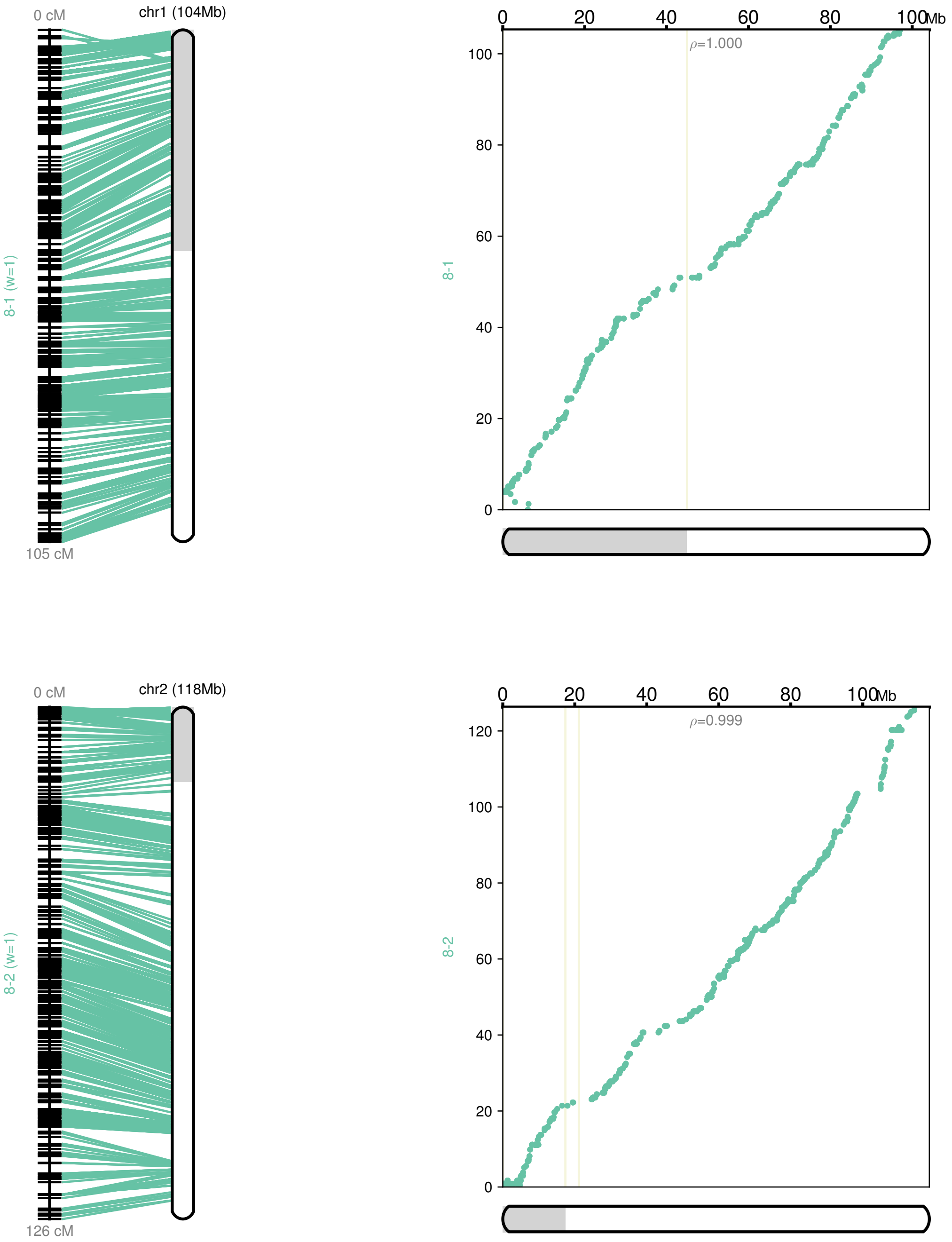


**Supplementary Figure S2** Genetic and physical maps of the acerola genome.

Left: SNP loci on the genetic map (vertical lines) and physical map (bars) are connected with horizontal lines. Right: Positions of SNP loci are indicated with dots on the genetic map (y-axis, cM) and physical map (x-axis, Mb).

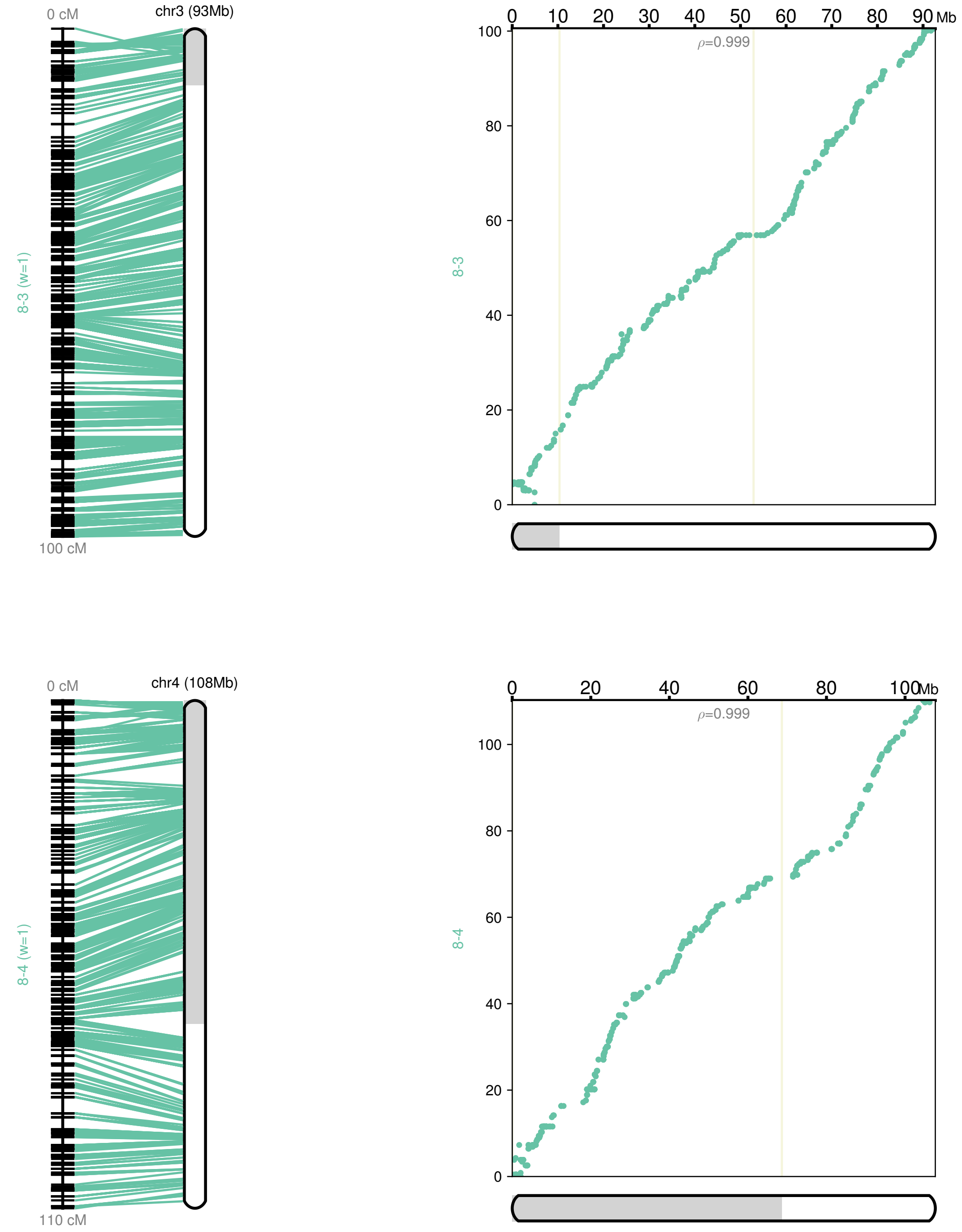


**Supplementary Figure S2** (continued)

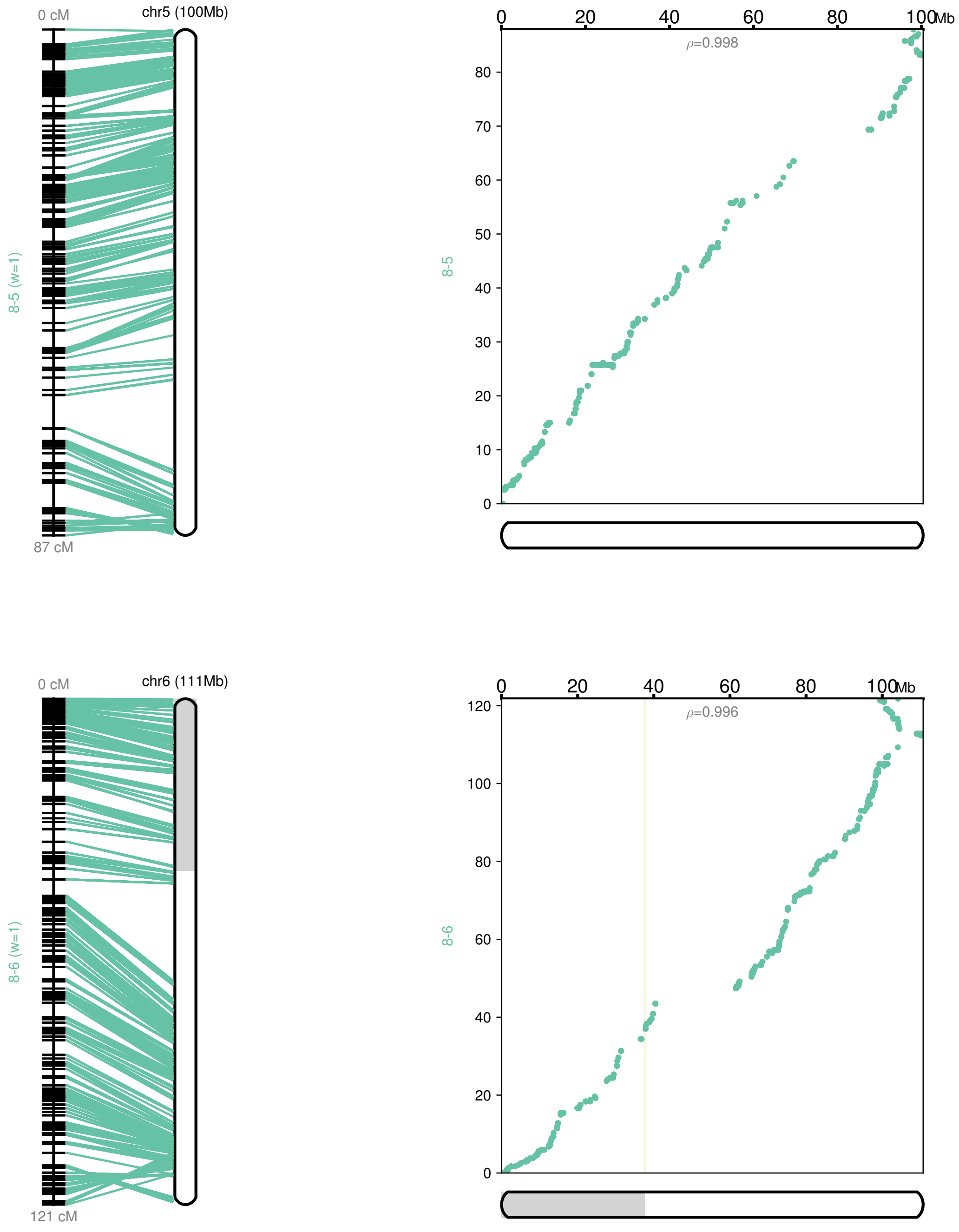


**Supplementary Figure S2** (continued)

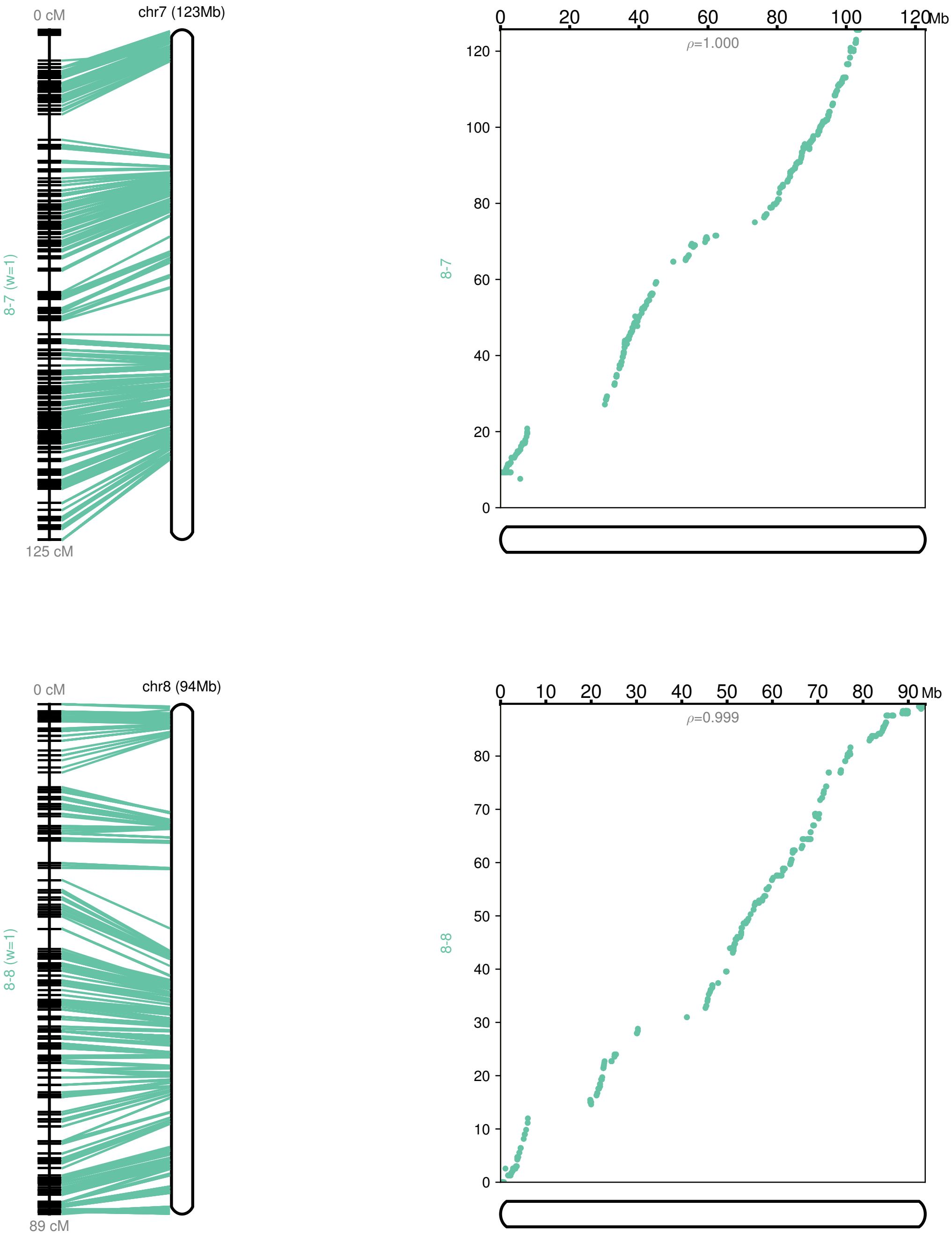


**Supplementary Figure S2** (continued)

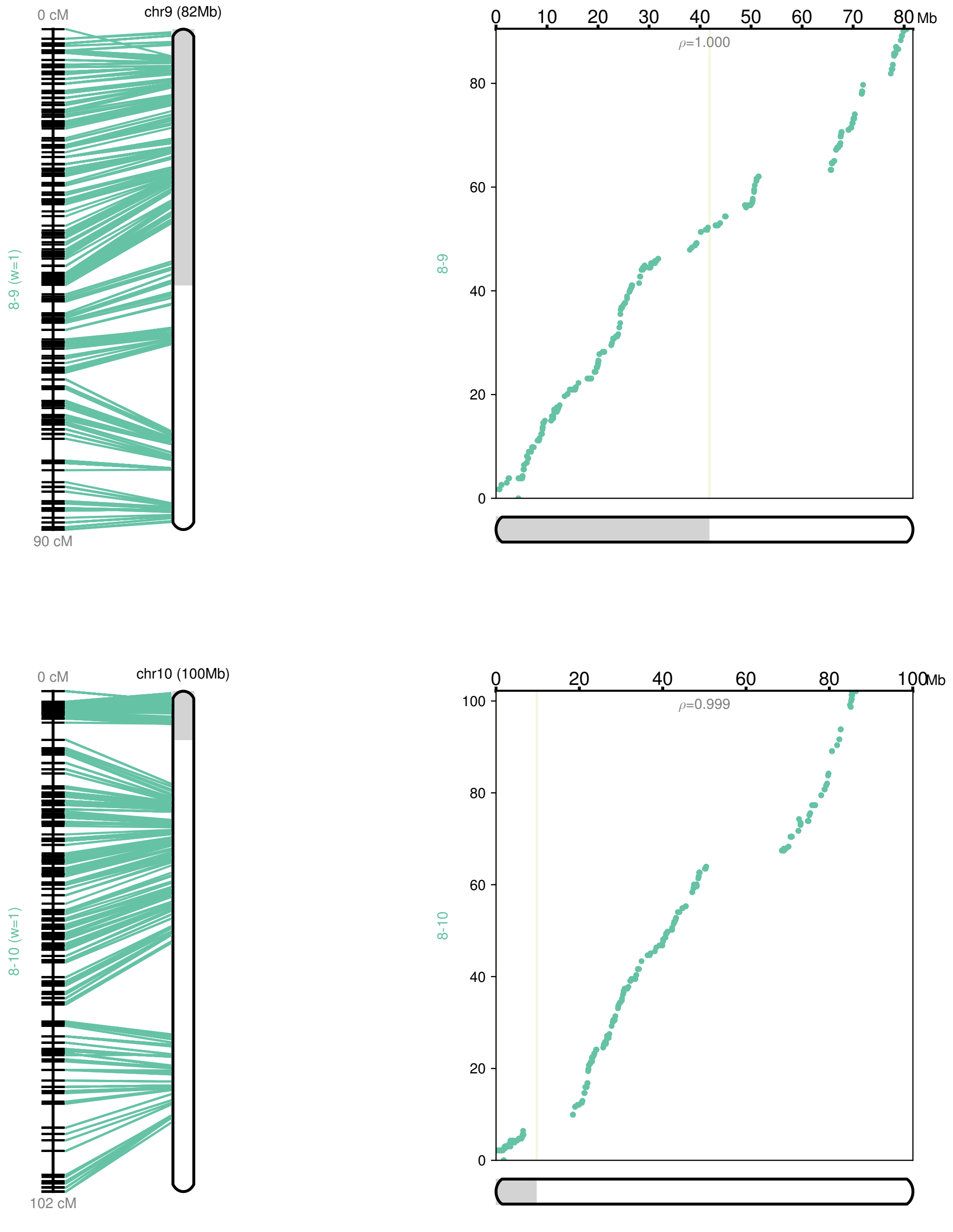


**Supplementary Figure S2** (continued)

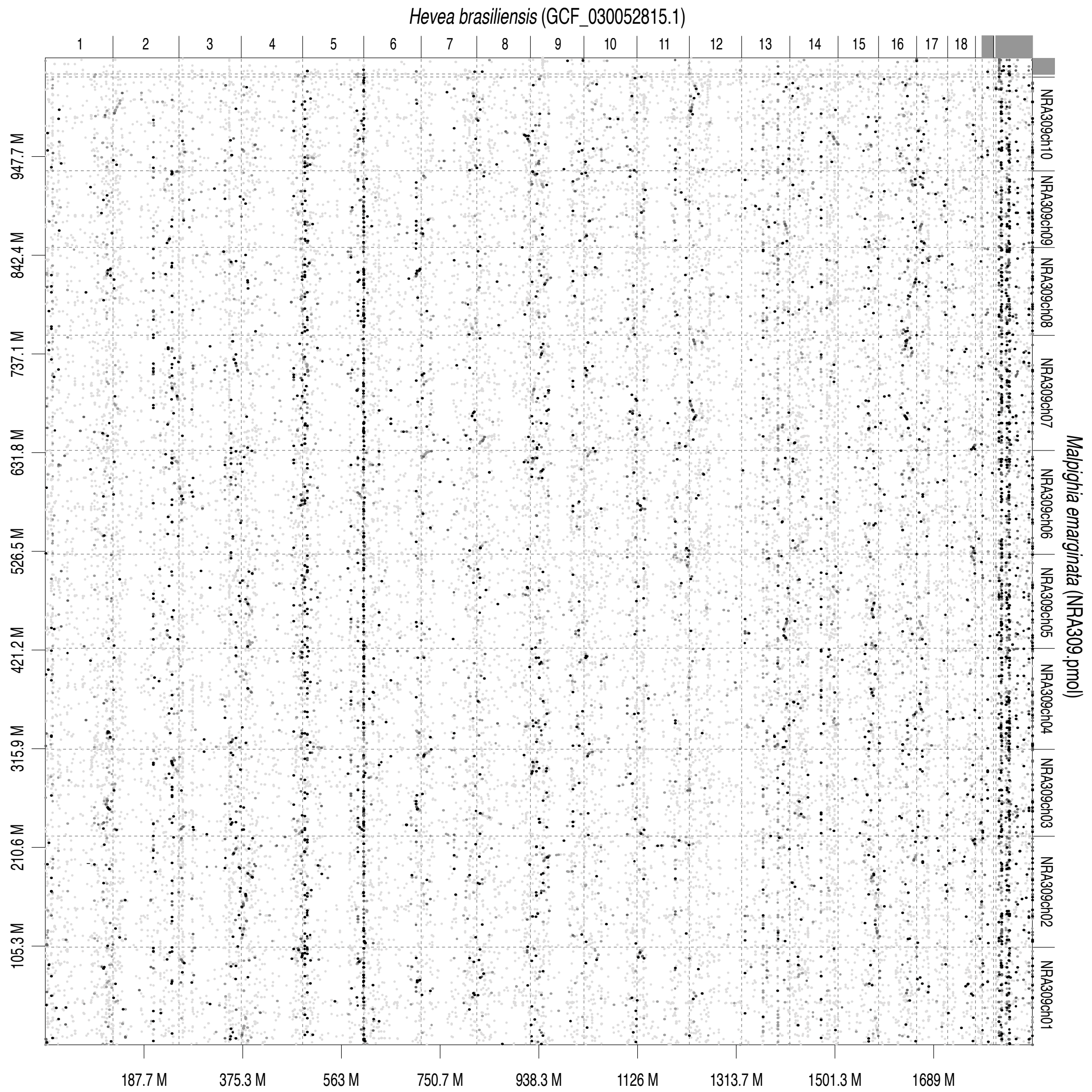


**Supplementary Figure S3** Comparative analysis of the genome sequence and structure of acerola and three other Malpighiales species.

Dots indicate similarities in the genome structures of rubber tree (*Hevea brasiliensis*), cassava (*Manihot esculenta*), and castor bean (*Ricinus communis*). Chromosome numbers are indicated above the x-axis and on the right side of the y-axis, and genome sizes (Mb) are shown below the x-axis and on the left side of the y-axis.

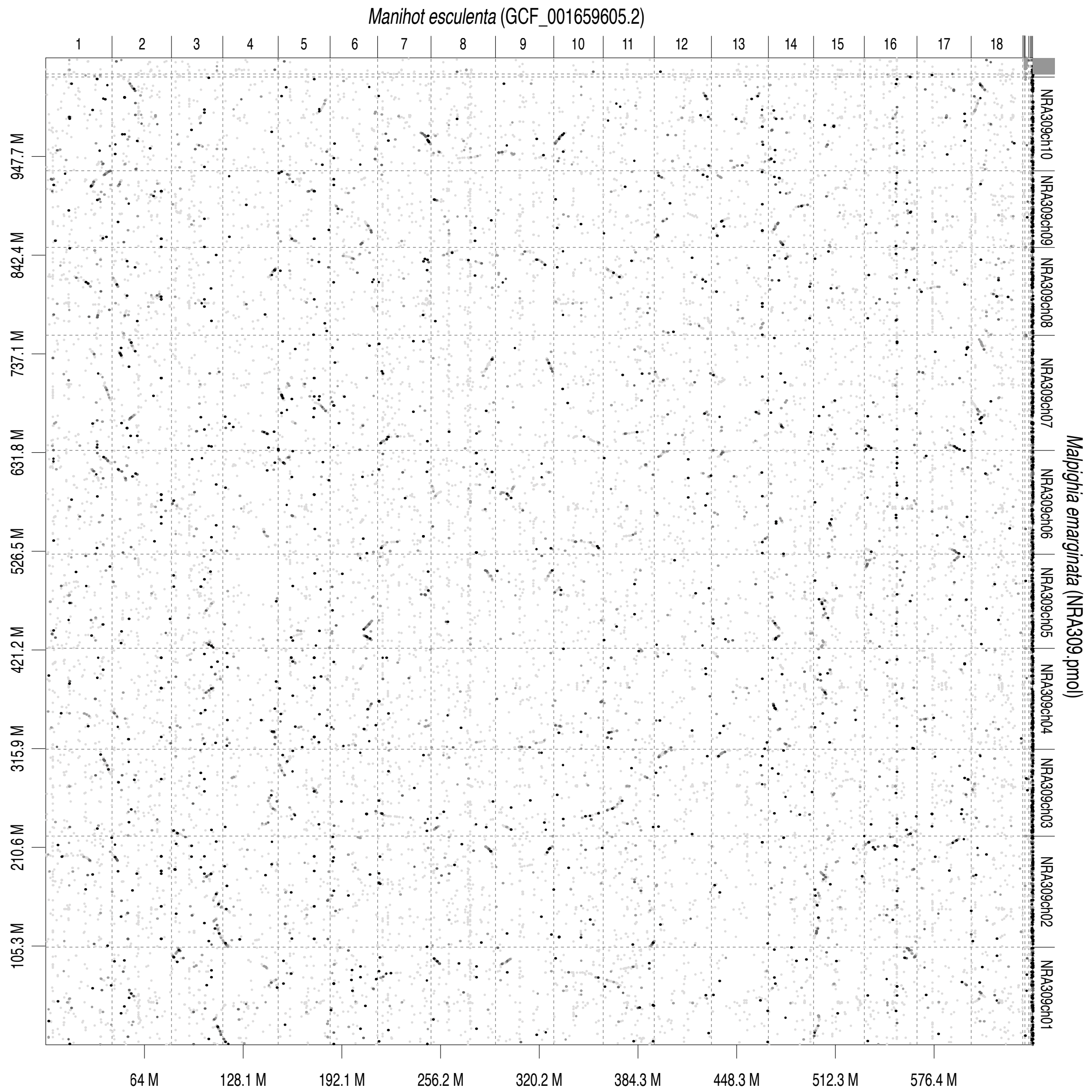


**Supplementary Figure S3** (continued)

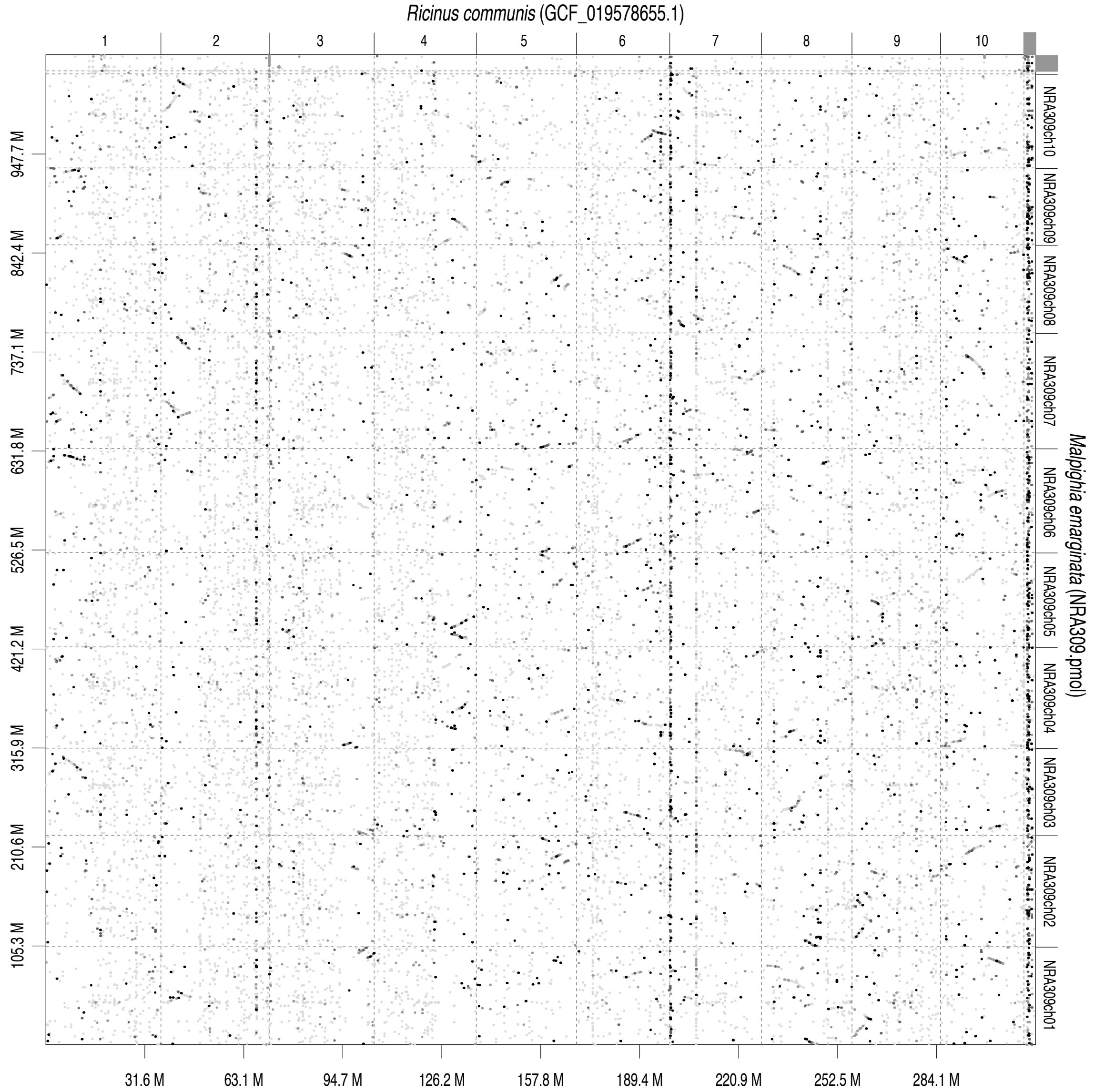


**Supplementary Figure S3** (continued)
